# Bright and sensitive red voltage indicators for imaging action potentials in brain slices and pancreatic islets

**DOI:** 10.1101/2022.12.01.518652

**Authors:** Yi Han, Junqi Yang, Yuan Li, Yu Chen, Huixia Ren, Ran Ding, Weiran Qian, Keyuan Ren, Beichen Xie, Mengying Deng, Yinghan Xiao, Jun Chu, Peng Zou

## Abstract

As fast developing tools for observing cellular membrane potential, red-emitting genetically encoded voltage indicators (GEVIs) reduce auto-fluorescence background, allow multiplexed recordings, and enable all-optical electrophysiology, but have been limited by either insensitivity or dimness. Here, we report a pair of red GEVIs, Cepheid1b/s, with improved sensitivity, brightness, and photostability. Cepheid1 indicators faithfully report cellular excitability in pancreatic islets and neural activity in acute brain slices.

## Main

Fluorescent indicators are powerful tools for spatiotemporally revolved mapping of cellular activities. Genetically encoded voltage indicators (GEVIs) allow non-invasive readout of membrane potential changes of large neuronal ensembles at the single-neuron level, thus enabling millisecond-timescale recording of neuronal activities, including subthreshold potentials^1^. A common challenge associated with voltage imaging is its high noise level due to limited photon counts, which arises from a combination of high acquisition frame rate (typically at 0.5-2 kHz) and low copy number of the membrane-embedded sensor protein. This high level of imaging noise has a significant impact on the signal-to-noise ratio (SNR), particularly for voltage imaging *in vivo*, which is further complicated with the presence of tissue auto-fluorescence and light scattering^2^. For this reason, GEVIs with red-shifted emission spectra are highly sought after, since they avoid much of the auto-fluorescence window and suffer less from scattering. Moreover, a brighter fluorophore could increase the baseline fluorescence signal which, in turn, promotes the SNR.

Existing red GEVIs are limited by speed, sensitivity, or brightness. For example, VSD-based GEVIs exhibit response time constants exceeding 5 ms^3, 4^; FlicRs^5, 6^ have fast kinetics but low sensitivity and dim fluorescence, which limited their performance in tissue. In comparison, rhodopsin-based GEVIs (e.g. QuasArs^7–9^ and Archons^10, 11^) feature sub-millisecond kinetics and high sensitivity (20% to 50% ΔF/F_0_ per AP); yet they suffer from low brightness due to limited quantum yield (<4%)^10^. To improve molecular brightness, a red fluorescent protein (RFP) donor is C-terminally fused to a voltage-sensing rhodopsin, which serves as the electrochromic Förster resonance energy transfer (eFRET) quencher. The resulting eFRET GEVIs, including QuasAr2-mRuby2^12^ and VARNAMs^6, 13^, exhibit substantially improved brightness while maintaining fast response kinetics. However, their absolute sensitivities are limited to ~20% ΔF/F_0_ per 100 mV^6, 12, 13^ owing to low FRET efficiencies^14, 15^.

Herein, we report a pair of red eFRET GEVIs with improved voltage sensitivity, brightness and photostability. Since voltage sensitivity critically depends on the FRET efficiency^15, 16^, we chose Ace^D81C^ with a red-shifted absorption spectrum^15^ as the voltage-sensing module to maximize the spectral overlap with RFP emission. Whereas previous eFRET GEVI designs have focused on C-terminally fused RFPs, our computational modeling (Figure S1) with AlphaFold2^17^ predicts 4-12 Å shorter distance and higher orientation factor (κ^2^) when RFP is inserted into the first extracellular loop (ECL1) of Ace rhodopsin (Figure 1a and Table S1).

**Figure 1.**
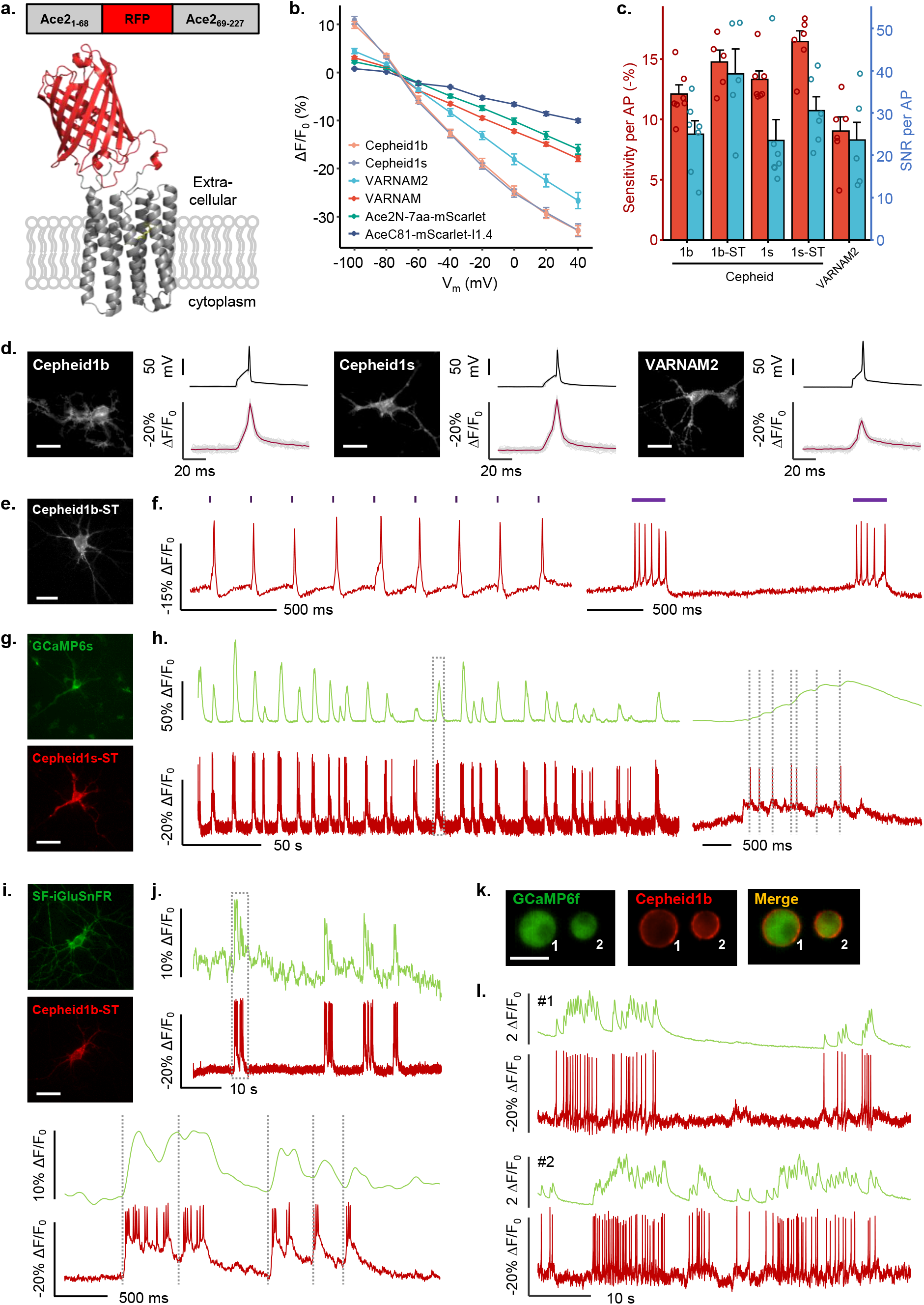
Design and Characterization of Cepheid1 indicators in cultured cells. **(a)** Diagram showing the RFP insertion site (top) and the predicted structure of Cepheid1s. **(b)** Normalized fluorescence-voltage response curve of red GEVIs. **(c)** Sensitivity of red GEVIs towards action potentials (APs) in cultured neurons. **(d)** Optical waveforms of Cepheids and VARNAM2 to APs. **(e-f)** Epifluorescence image (**e**) and whole-cell fluorescence response (**f**) of cultured neuron expressing Cepheid1b-ST-P2A-CheRiff, when optogenetically triggered with 405 nm laser. **(g-h)** Epifluorescence images (**g**) and whole-cell fluorescence traces (**h**) of cultured rat hippocampal neuron expressing GCaMP6s-NES (top) and Cepheid1s-ST (bottom), with zoom-in view of the boxed region shown on the right. **(i-j)** Epifluorescence images (**i**) and whole-cell fluorescence traces (**j**) of cultured rat hippocampal neuron expressing SF-iGluSnFr (top) and Cepheid1b-ST (bottom), with zoom-in view of the boxed region shown at the bottom. **(k-l)** Epifluorescence images (**k**) and whole-cell fluorescence traces (**l**) of mouse pancreatic islet cells expressing GCaMP6f (cyan) and Cepheid1b (red). Scale bars, 20 μm.

Our list of RFP candidates include mRuby3^18^, mRuby4^37^, mScarlet^19^. We also included an engineered variant of mScarlet-I, mScarlet1.4 (unpublished), which is spectrally similar (Figure S1 and Table S2) but with better folding (Figure S2) and higher brightness (Figure S3). To improve membrane trafficking, a combination of triple-tandem transport signal (3xTS) and ER-exiting (ER2) sequences^20^ are C-terminally fused to the rhodopsin (Figure 1a). Except for mScarlet, all RFP-Ace insertions exhibited good expression and membrane trafficking in human embryonic kidney 293T (HEK293T) cells (Figure S4). While mScarlet1.4 insertion showed the brightest fluorescence (Figure S5), mRuby4 insertion exhibited superior photostability, with half-bleach time more than 13 min when illuminated with 561 nm laser at 1.6 W/cm^2^ (Figure S6). We therefore named mScarlet1.4- and mRuby4-insertions as Cepheid1b (for brighter fluorescence) and Cepheid1s (for better photostability), respectively.

We then applied whole-cell voltage clamping in HEK293T cells to characterize the response kinetics and dynamic range of Cepheid1 indicators. Upon membrane depolarization from −70 mV to 30 mV, both indicators exhibited millisecond kinetics with 1.9 ± 0.1 ms and 1.5 ± 0.2 ms (Figure S7). The voltage sensitivity of Cepheid1b and Cepheid1s compare favorably against several recently published red GEVIs, including VARNAM^6^, VARNAM2^13^, and Ace2N-7aa-mScarlet^21^, which are constructed by fusing mRuby3 and mScarlet to the C-terminus of Ace rhodopsin mutants. When the membrane potential is altered step-wise from −100 mV to +40 mV (Figure S8), the whole-cell fluorescence of VARNAM, VARNAM2, and Ace2N-7aa-mScarlet change by −17.9 ± 0.6%, −26.7 ± 1.7% and −16.0 ± 1.0% ΔF/F_0_, respectively (Figure 1b). In comparison, Cepheid1b (−32.9 ± 1.0%) and Cepheid1s (−33.6 ± 1.5%) are nearly 2-fold as sensitive as VARNAM and Ace2N-7aa-mScarlet, and ~23% higher than VARNAM2. Notably, fusing mScarlet1.4 to the C-terminus of Ace^C81^ reduces the voltage sensitivity by more than 2-fold, which supports the computational modeling prediction (Figure 1b).

We applied Cepheid1 indicators to image action potentials (APs) in cultured rat hippocampal neurons. Both indicators exhibited good expression and membrane localization in soma and neurites (Figure 1d). The sensitivity of Cepheid1b (−12.1 ± 0.8%) and Cepheid1s (−13.3 ± 0.7%) towards APs (ΔF/F_0_) are more than 34% higher than VARNAM2 (9.0 ± 1.2%) (Figure 1c). To improve SNR, we fused Cepheid1 indicators with a 63-amino acid somatic targeting (ST) domain from potassium channel Kv2.1, which restricts their expression to the soma and proximal dendrites^22^. The resulting soma-targeted Cepheid1b/s-ST exhibit 40% improved SNR (Figure 1c).

The red-shifted spectra of Cepheid1 indicators allow the combination with the photosensitive cation channel CheRiff^7^ for all-optical electrophysiology. In cultured rat hippocampal neurons co-expressing CheRiff and Cepheid1b/s-ST, we applied 405 nm laser to stimulate AP firing while simultaneously imaging membrane voltage (Figures 1e and S9). Under various stimulation patterns, Cepheid1b-ST faithfully recorded APs (Figure 1f), while Cepheid1s-ST allows for long-term imaging over 20 min (Figure S10).

We further paired Cepheid1b/s-ST with green-emitting indicators to simultaneously imaging membrane potential with other neuronal signals such as cytoplasmic calcium and extracellular glutamate. We co-expressed Cepheid1b/s-ST with GCaMP6s^23^ in primary rat hippocampal neurons (Figures 1g and S11). The high photostability of Cepheid1s-ST allowed continuous imaging of time-correlated voltage and calcium spikes for 5 min (Figures 1h). We also applied Cepheid1b-ST and SF-iGluSnFR^24^ to record voltage-induced glutamate release (Figure 1i-j and Figure S12). In addition, the high brightness of Cepheid1b and its spectral orthogonality with GCaMP6f have enabled us to sensitively record membrane potential and cytoplasmic calcium in cultured mouse pancreatic islet cells, which unveils time-correlated spontaneous calcium waves and AP spikes that are reminiscent to neuronal activities (Figure 1k-l).

To assess Cepheid1’s performance in tissue, we performed voltage imaging in mouse brain slices and pancreatic islets. We introduced Cepheid1b-ST into mouse brain via adeno-associated virus infection, which showed good expression and membrane trafficking across multiple brain regions, including cortex, hippocampus, and cerebellum (Figure 2a). In acute brain slices, the high sensitivity of Cepheid1b-ST (ΔF/F_0_ = - 9.9 ± 0.3 % per AP, Figure 2b) has allowed faithfully report firing APs in single trial measurements (Figure 2c) at the frame rate of 498 Hz.

**Figure 2.**
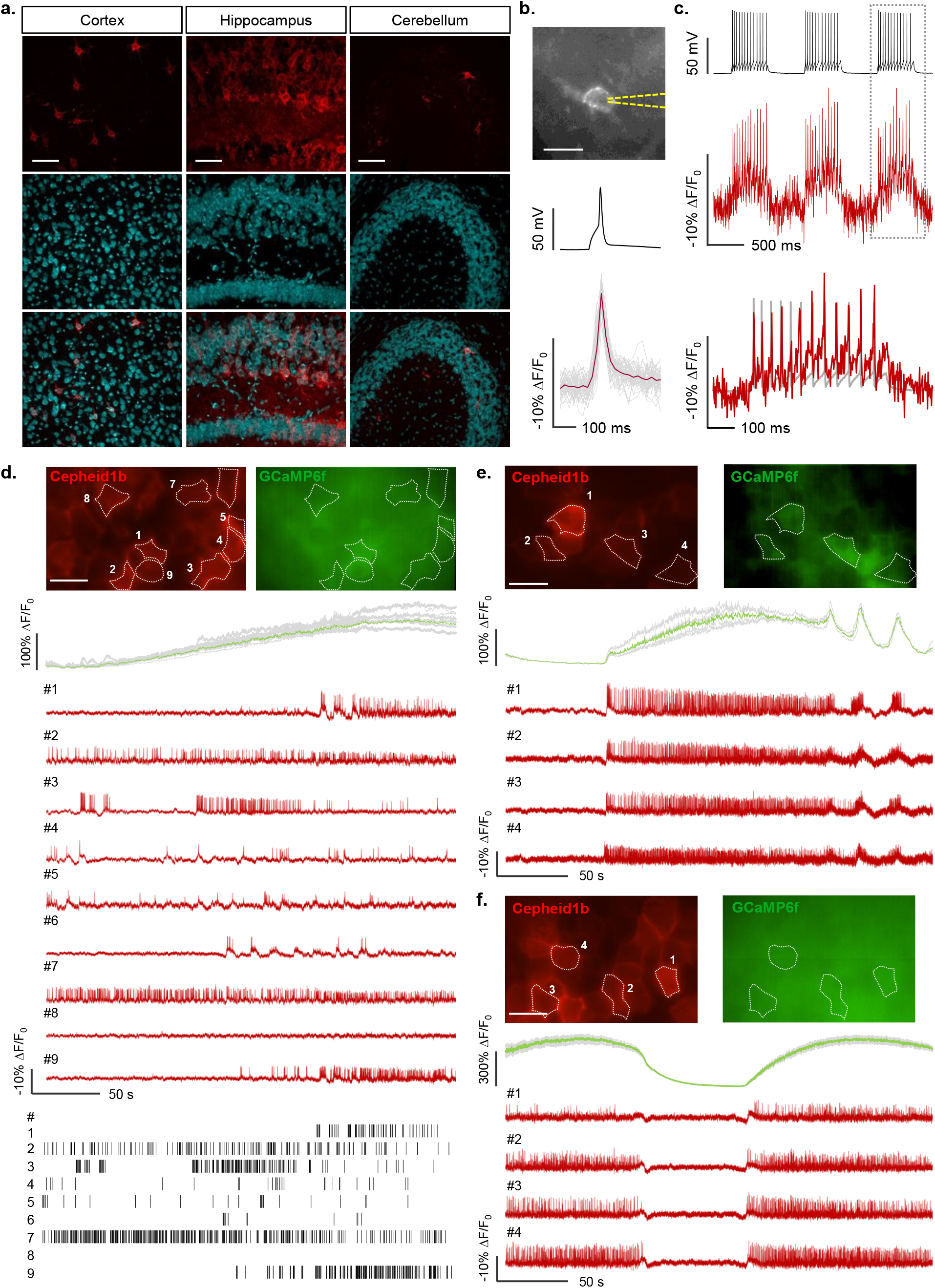
Voltage imaging with Cepheid1 indicators in acute brain slice and pancreatic islets *ex vivo*. **(a)** Confocal images of fixed slices showing expression and localization of Cepheid1b-ST (scale bar = 50 μm). **(b)** Fluorescence image (top), mean electrical (black) and optical (red) waveforms of stimulated APs recorded from a hippocampal neuron expressing Cepheid1b-ST in acute brain slice. Grey and red lines indicate individual and mean optical traces, respectively (*n* = 48 spikes). **(c)** Fluorescence response (red) of Cepheid1b-ST to single-trial burst firing (black) in acute brain slice, with zoom-in view of the boxed region shown at the bottom. **(d-f)** Representative dual-color imaging of calcium (mean, cyan; individual, gray) and voltage (red) in isolated islets at 200 Hz frame rate. At the onset of imaging, extracellular glucose is switched from 3 mM to 10 mM (**d**). Following 30-60 mins incubation at 10 mM glucose, both mixed (**e**) and slow (**f**) calcium oscillations are observed together with time-correlated electrical spikes. Scale bars, 20 μm.

Glucose level-dependent calcium oscillations have been observed in mammalian pancreatic islets^25^, yet the interplay between membrane potential and intracellular calcium has not been investigated. We thus infected *GCaMP6f*^+/+^ mice with adenovirus encoding Cepheid1b and performed dual-color imaging in isolated pancreatic islets. Upon switching from low (3 mM) to high (10 mM) levels of extracellular glucose, we observed highly heterogenous patterns of electrical activities in individual islet cells, whereas their calcium signal increased gradually and synchronously over the time course of minutes (Figure 2d). Interestingly, following 30-60 mins incubation in high-glucose medium, the electrical activities of islet cells appear highly synchronized and time-correlated with both mixed and slow calcium oscillations (Figure 2e-f).

To summarize, we have reported a pair of red GEVIs with improved voltage sensitivity of 33% −ΔF/F_0_ per 100 mV, which is the highest among red GEVIs reported to date. The high dynamic range of Cepheid1 allows sensitive detection of APs at 500 Hz frame rate in both cultured neurons (ΔF/F_0_ = −12%) and brain slices (ΔF/F_0_ = −10%) under mild illumination with 561 nm laser lower than 2 W/cm^2^. While Cepheid1b are useful for detecting both subthreshold and action potentials as a brighter indicator, Cepheid1s supports long-term imaging due to its high photostability. Both GEVIs support multiplexed imaging with green fluorescent indicators and all-optical electrophysiology. We further demonstrated that Cepheid1b faithfully report spike firing in acute mouse brain slices and pancreatic islets. Unlike previous engineering efforts that focused exclusively on C-terminal RFP fusions, we achieved higher voltage sensitivity through insertion into ECL1. Such novel design leveraged the predictive power of computational modeling of chimeric protein structures. We envision that the sensitivity of Cepheid1 indicators could be further optimized through directed evolution.

## Methods

### Materials and reagents

The reagents used in this study are summarized in Table S3.

### Molecular cloning

Plasmids used in this study were generated by Gibson Assembly ligating PCR amplified inserted DNA fragment and linearized vector with 25-bp overlap. The primers for PCR are summarized in Table S4. DNA fragments and linearized vector were mixed with Gibson Assembly enzyme (Lightening Cloning Kit). The sequences of plasmids were verified through Sanger sequencing.

### Computational modeling of FRET efficiency

The structures of Cepheid1b and Cepheid1s are predicted with AlphaFold2. Non-amino-acid chromophores are manually added via structural alignment of the predicted structures against the crystallography data of separate protein domains: Acetabularia Rhodopsin II (PDB ID: 3AM6), mScarlet (PDB ID: 5LK4) and mRuby (PDB ID: 3U0L) as proxies of mScarlet1.4 and mRuby4, respectively. The coordinates of fluorophores are used for calculating the distance R (defined as the distance between the centers of mass of donor and acceptor) and the orientation factor *κ*^2^ (defined as 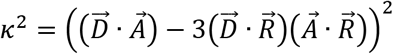, where 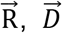 and 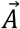 are vectors connecting the centers of mass of donor and acceptor, normalized longitudinal axis of donor and acceptor, respectively.

### Expression of Cepheid1 in cultured cells

HEK293T cells were seeded in a 24-well plate and incubated in Dulbecco’s modified Eagle medium (EMEM, Gibco) containing 10% v/v fetal bovine serum (FBS, Gibco) at 37 °C with 5% CO_2_. Cells were transfected with Lipofectamine 3000 reagent following the manufacturer’s instructions. Primary rat hippocampal neurons were digested from dissected rat brain at postnatal day 0 (P0) and plated onto sterile 14-mm glass coverslips pre-coated with 20 μg/mL poly-D-lysine and 10 μg/mL laminin. Neurons were incubated at 37 °C with 5% CO_2_ and were transfected on DIV7-9 (7-9 days *in vitro*) with Lipofectamine 3000 following the manufacturer’s instructions. Transfected neurons were imaged after 3-7 days.

### Imaging apparatus

Fluorescence microscopy in cultured cells and pancreatic islets was performed on an inverted microscope (Nikon-TiE) equipped with a 40x, NA1.3 oil immersion objective lens, three laser lines (Coherent OBIS 405 nm, 488 nm and 561 nm), a spinning disk confocal unit (Yokogawa CSU-X1) and two scientific CMOS cameras (Hamamatsu ORCA-Flash 4.0 v2). A dual-view device (Photometrics DV2) was used to split the emission into green/red fluorescence channels. Fluorescence imaging experiments in acute slices were performed on an upright microscope (Olympus-BX51WI) equipped with a 40x, NA0.8 water immersion objective lens, a 561 nm laser line (Coherent OBIS) and a scientific CMOS camera (Hamamatsu ORCA-Flash 4.0 v2). The spectra properties of the filters and dichroic mirrors for various fluorescent indicators used in this study are summarized in Table S5.

### Fluorescence voltage imaging in HEK293T cells and cultured neurons

To measure the dynamic range and kinetics of the voltage indicators in HEK293T cells, the membrane potential was controlled via whole-cell patch clamp (Axopatch 200B, Axon Instruments). Fluorescence imaging was performed at the frame rate of 1058 Hz. To measure the response of indicators to action potentials, neurons were current-clamped and injected with 200-500 pA of current for 5-10 ms to stimulate action potential firing. Fluorescence imaging was performed at the frame rate of 484 Hz. For simultaneous imaging of membrane voltage and calcium, Cepheid1 and GCaMP6s-NES were co-transfected into neurons at DIV8. On DIV14-18, neurons were illuminated with 488 nm (2.4 W/cm^2^) and 561 nm (1.6 W/cm^2^) lasers and imaged with a dual-view device at a frame rate of 500 Hz. For simultaneous imaging of membrane voltage and glutamate, Cepheid1 and iGluSnFR were co-transfected into neurons on DIV8 and imaged on DIV14-19 in dual-view mode (2.4 W/cm^2^ of 488 nm and 1.6 W/cm^2^ of 561 nm laser) at a frame rate of 500 Hz.

### Electrophysiology for cultured cells

Borosilicate glass capillaries (BF150-86-10, Sutter) were pulled with a micropipette puller (P-1000, Sutter) and the pipette resistance was 4–8 MΩ. The extracellular solution for whole-cell patch clamp contains 125 mM NaCl, 2.5 mM KCl, 15 mM HEPES and 30 mM glucose (305-310 mOsm/kg, pH 7.3). The intracellular solution contains 125 mM potassium gluconate, 8 mM NaCl, 1 mM EGTA, 10 mM HEPES, 0.6 mM MgCl_2_, 0.1 mM CaCl_2_ (pH 7.3, 295 mOsm/kg). Whole-cell current was recorded with an Axopatch 200B amplifier (Axon Instruments). Membrane voltage signal recorded from the patch amplifier was filtered with an internal 5-kHz Bessel filter and digitized at 9,681.48 Hz with a National Instruments PCIe-6353 data acquisition (DAQ) board (approximately twice the bandwidth of the Bessel filter).

### All-optical electrophysiology

Cultured hippocampal neurons were transfected with plasmid encoding CheRiff and soma-targeting Cepheid1b/s (Cepheid1b/s-ST) mediated by P2A self-cleaving peptide. Transfected neurons were activated with repeated 2 ms (0.5 W/cm^2^) 405 nm light pulses, or repeated 250-300 ms (0.1-0.3 W/cm^2^) 405 nm light steps, or gradually enhanced (from 0 to 0.2 W/cm^2^ in 10 s) 405 nm light. Images were captured at a frame rate of 500 Hz under constant 561 nm illumination (1.6 W/cm^2^). For each trial, 25 μM APV, 10 μM NBQX and 20 μM gabazine were added into extracellular solution to block synaptic transmission.

### Intracerebroventricular injection and acute slice measurements

C57BL/6N mouse lines were purchased from Charles River. The AAV vector expressing Cepheid1b-ST: AAV2/PHP.eb-hsyn-Cepheid1b-ST was custom-produced by Chinese Institute for Brain Research (CIBR). For one 6 to 7-week-old mouse (without regard to sex), 3 μL of AAV2/PHP.eb-hsyn-Cepheid1b-ST (4.8 x 10^12^ GC mL^-1^) was injected into the lateral ventricle. The coordinate for intracerebroventricular injection (in mm from Bregma: AP, ML) was −0.58 – −0.59, 1.35 – 1.4, DV = 1.8 – 1.9 mm.

Acute slices were prepared from 9 to 11-week-old mice (at least 3 weeks after AAV injection). Mouse was deeply anesthetized via isoflurane inhalation and rapidly decapitated. The brain was dissected from the skull and placed in ice-cold artificial cerebrospinal fluid (ACSF) containing 26 mM NaHCO_3_, 1.25 mM NaH_2_PO_4_, 125 mM NaCl, 2.5 mM KCl, 2 mM CaCl_2_, 1 mM MgCl_2_, 2 mM KCl and 20 mM glucose (pH 7.3-7.4, 295 mOsm/kg) and saturated with Carbogen (95% O_2_ and 5% CO_2_). The brain was sliced into 250 μm sections with a Leica VT1200s vibratome. Slices were incubated for 45 min at 34.5 °C in ACSF and maintained at room temperature (22 °C). ACSF was continuously bubbled with Carbogen for the duration of the preparation and subsequent experiment.

For fluorescence imaging and electrophysiology recoding, slices were placed on a custom-built chamber and held by a platinum harp net stretched across. Carbogen-bubbled ACSF was perfused at a rate of 4 mL/min with a Longer peristaltic pump. Fluorescence images were captured at a frame rate of 400-500 Hz, under 2 W/cm^2^ 561 nm illumination. The intracellular solution for whole-cell patch clamp contains 105 mM potassium gluconate, 30 mM KCl, 4 mM MgATP, 0.3 mM Na_2_-GTP, 0.3 mM EGTA, 10 mM HEPES and 10 mM sodium phosphocreatine (pH 7.3, 295 mOsm/kg). Electrophysiology measurements were acquired with a MultiClamp 700B amplifier (Molecular Devices).

### Isolation of mouse pancreatic islets and infection of Cepheid1b

Islets of Langerhans were isolated from *GCaMP6f*+/+ mice. After isolation, the islets were cultured overnight in RPMI-1640 medium containing 10% fetal bovine serum, 8 mM D-glucose, 100 unit/ml penicillin and 100 mg/mL streptomycin for overnight culture at 37 °C in a 5% CO_2_ humidified air atmosphere. The islets were infected with adenoviruses pAdeno-CMV-Cepheid1b by 1.5 h exposure in 100 μL culture medium, with approximately 4×10^9^ plaque forming units (PFU) per islet, followed by addition of regular medium and further culture for 16-20 h before use.

### Dissociation into islet single cells

Freshly isolated islets were washed with HBSS and subsequently digested with 0.025% trypsin-EDTA for 3 min at 37 °C followed by brief shaking. Digestion was stopped with addition of culture medium, and the solution was centrifuged at 94 g for 5 min. The cells were suspended by RPMI 1640 culture medium. The cell suspension was plated on coverslips in the poly-L-lysine-coated Glass Bottom Dish (D35-14-1-N, Cellvis) or a microfluidic chip. The dishes or chips were then kept for 60 min in the culture incubator at 37 °C, 5% CO_2_ to allow cells to adhere. Additional culture medium was then added, and the cells were cultured for 24 h before the imaging experiments.

### Voltage imaging in mouse pancreatic islets and islet single cells

For simultaneous imaging with Cepheid1b and GCaMP6f, mouse pancreatic islets and mouse islet single cells were illuminated with 488 nm and 561nm lasers at 0.2–0.5 W/cm^2^ and 0.9–1.8 W/cm^2^, respectively, and continuously imaged for 60–300 s at a camera frame rate of 200 Hz. The samples were kept at 37 °C on the microscope stage during imaging. For mouse islets, a polydimethylsiloxane (PDMS) microfluidic chip was used to provide a stable and controllable environment for long-term imaging. The reagents were automatically pumped into the microfluidic chip with a flow rate of 800 μL/h by the TS-1B syringe pump (LongerPump). Before imaging, the chip and all the solution were degassed with a vacuum pump for 5 min to achieve stable hour-long imaging. The microfluidic chip was pre-filled with KRBB solution (125 mM NaCl, 5.9 mM KCl, 2.56 mM CaCl_2_, 1.2 mM MgCl_2_, 1 mM L-glutamine, 25 mM HEPES, 0.1% BSA, pH 7.4) containing 3 mM D-glucose before use.

### Data analysis

Most fluorescence images and electrophysiology recordings were analyzed with home-built software written in MATLAB (MathWorks, version R2022a). The fluorescence images obtained in acute slices were pre-processed with a python-based voltage imaging data analysis package (VolPy). Fluorescence intensities were extracted from the mean values over a manually drawn region of interest around the soma of each labelled cell. Statistical analysis was performed with Origin (version 2019b) and R (version 4.2.0).

## Acknowledgments

This work was supported by the Ministry of Science and Technology (2018YFA0507600, 2017YFA0503600), the National Natural Science Foundation of China (32088101, 21727806). PZ is sponsored by Bayer Investigator Award.

## Author Contributions

J.Y. and P.Z. conceived the project. Y.H., JY., Y.L., Y.C. and P.Z. designed experiments. Y.H., J.Y., Y.L., Y.C., H.R. K.R., B.X., M.D. and Y.X. performed experiments. Y.H., J.Y. and P.Z. analyzed data and wrote the paper with inputs from other authors.

## Declaration of Interests

The authors declare no competing interests.

## Notes

### Competing Interest Statement

The authors have declared no competing interest.

