## Supplemental Information for "Bright and sensitive red voltage indicators for imaging action potentials in brain slices and pancreatic islets"

### Supplementary Tables

**Table S1**: FRET efficiency predictions

| | **R (Å)** | **$\kappa^2$** | **$\kappa^2/R^6$ ($10^{-11}$ Å$^{-6}$)** |
|---|---|---|---|
| $Ace^{81C}$-(C)mScarlet-I1.4 | 63 | 0.00026 | 0.00042 |
| $Ace^{81C}$-(C)mRuby4 | 52 | 0.0045 | 0.023 |
| $Ace^{81C}$-(ECL1)mScarlet-I1.4 | 51 | 0.26 | 1.5 |
| $Ace^{81C}$-(ECL1)mRuby4 | 48 | 0.66 | 5.4 |

**Table S2**: Photophysical properties of mScarlet-I1.4 compared with mScarlet-I

| | mScarlet-I | mScarlet-I1.4 |
|---|---|---|
| Absorption maximum | 570 nm | 570 nm |
| Excitation maximum | 571 nm | 571 nm |
| Emission maximum | 511 nm & 600 nm | 512 nm & 600 nm |
| Green/Red emission ratio | 0.502 | 0.119 |
| Relative brightness in bacteria | 0.489 / 0.301 | 1 |

**Table S3**: List of reagents used in this study

| **Reagent** | **Vendor** | **Catalog Number** |
|---|---|---|
| Phanta® Max Super-Fidelity DNA Polymerase | Vazyme | P505-d2 |
| Lightening Cloning Kit | Biodragon | BDIT0014-100 |
| DNA extraction kit | TIANGEN | DP118-02 |
| Dulbecco's Modified Eagle's medium (DMEM) | Gibco | C11995500BT |
| Fetal Bovine Serum (FBS) | Gibco | 10099141C |
| Trypsin-EDTA (0.25%) | Gibco | 25200056 |
| Neurobasal™ Medium | Gibco | 21103049 |
| B-27™ Supplement | Gibco | 17504044 |
| GlutaMAX™ Supplement | Gibco | 35050061 |
| Penicillin-streptomycin | Beyotime | C0222 |
| Matrigel® Matrix | Corning | 356234 |
| poly-D-lysine | Sigma | P7280-5X5M |
| Laminin Mouse Protein | Gibco | 23017015 |
| Opti-MEM® Medium | Gibco | 31985062 |
| Lipofectamine® 3000 Reagent | Gibco | L3000008 |
| HEPES | Amresco | 0511 |
| EGTA | Sigma | 03777-10G |
| 2-APB | Abcam | ab120124 |
| Gabazine | Abcam | ab120042 |
| NBQX | Abcam | ab120045 |
| D-AP5 (APV) | Abcam | ab120003 |
| Creatine Phosphate Sodium | Maklin | C804629 |
| Potassium gluconate | Sigma | G4500 |
| Adenosine 5'-triphosphate Magnesium | Maklin | A922363 |
| HBSS | Gibco | C14175500BT |
| $NaHCO_3$ | Tong Guang | 144-55-8 |

| | | |
|---|---|---|
| $NaH_2PO_4 \cdot 2H_2O$ | Xilong Scientific | 13472-35-0 |
| NaCl | Sigma | S3014 |
| Glucose | Sigma | G7021 |
| KCl | Sigma | P9541 |
| $MgCl_2$ | Sigma | M2670 |
| $CaCl_2$ | Sigma | 10043-52-4 |

**Table S4**: List of cloning primers used in this study

| Construct | Primer sequence (5' to 3') |
|---|---|
| Ace (N terminus) | Forward: ATGGCTGACGTGGAAACCGAG<br>Reverse: CATTGTCAGGTCCTGGTAGTTCACG |
| Ace (C terminus) | Forward: AATGGTGAAAGGCAGGTGGTCTAC<br>Reverse: CTTGAAGATAGTCTCATGGGCAATGAGG |
| mScarlet-I1.4 | Forward: TACCAGGACCTGACAATGATGGTGAGCAAGGGCGAGG<br>Reverse: CCTGCCTTTCACCATTCTTGTACAGCTCGTCCATGCCG |
| mRuby4 | Forward: CGTGAACTACCAGGACCTGACAATGCTGATCAAGGAAGAGATGCCCATGAAG<br>Reverse: CGTAGACCACCTGCCTTTCACCATTCTTGTACAGCTCGTCCATGCC |
| Δ7mOrange2(Y71F)-ER2 | Forward: CATCAATGTGGGGGGCAACATGGCCATCATCAAGGAGTTCATG<br>Reverse: AGCTTGATATCGAATTCTCATTACACCTCGTTCTCGTAGCAGAACTTGTA |
| 3xTS | Forward: CCCATGAGACTATCTTCAAGACCGGTGCCGCCGACCG<br>Reverse: GCCCCCCACATTGATGTCAATCTG |
| Kv2.1 motif | Forward: TGGACGAGCTGTACAAGCAGAGCCAGCCTATCCTGAACAC<br>Reverse: CGATAAGCTTGATATCGAATTCTTACACTTCATTTTCATAGCAGAAGAACCTGG |
| P2A-CheRiff | Forward: GGCTCCGGAGCCACGAAC<br>Reverse: AGCGTAATCTGGAACATCGTATGGG |

**Table S5**: Spectral properties and imaging apparatus for fluorescent imaging

| Indicator | Fluorophore | Excitation max. (nm) | Emission max. (nm) | Laser line (nm) | Emission filter (nm) |
|---|---|---|---|---|---|
| Cepheid1b | mScarlet-I1.4 | 570 | 600 | 561 | 630 / 75 (inverted wide-field)<br>600 / 50 (upright wide-field, confocal, dual-color) |
| Cepheid1s | mRuby4 | 558 | 592 | 561 | 630 / 75 (wide-field)<br>600/50 (confocal, dual-color) |
| VARNAM[1] | mRuby3 | 558 | 592 | 561 | 630 / 75 |
| VARNAM2[2] | mRuby3 | 558 | 592 | 561 | 630 / 75 |
| Ace2N-7aa-mScarlet[3] | mScarlet | 569 | 594 | 561 | 630 / 75 |
| AceC81-mScarlet-I1.4 | mScarlet-I1.4 | 570 | 600 | 561 | 630 / 75 |
| GCaMP6s[4] | cpEGFP | 497 ($Ca^{2+}$-bound) | 515 ($Ca^{2+}$-bound) | 488 | 525 / 50 |

Dichroic mirror: Chroma ZT405/488/561/640rpc for confocal imaging and ZT405/488/532/642rpc for inverted wide-field imaging. ZT561rdc for upright wide-field wide-field imaging. T565LPXR was used to split fluorescence for dual-color imaging.

### Supplementary Figures

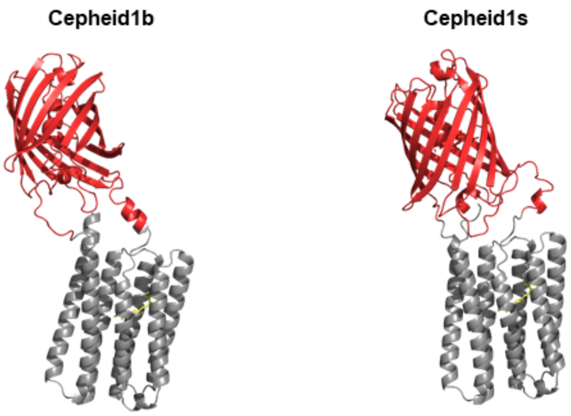


**Figure S1 Tertiary structure prediction of Cepheid1b and Cepheid1s by AlphaFold2.** Retinal and RFP fluorophore are added by manual alignment with crystallography data in RCSB PDB.

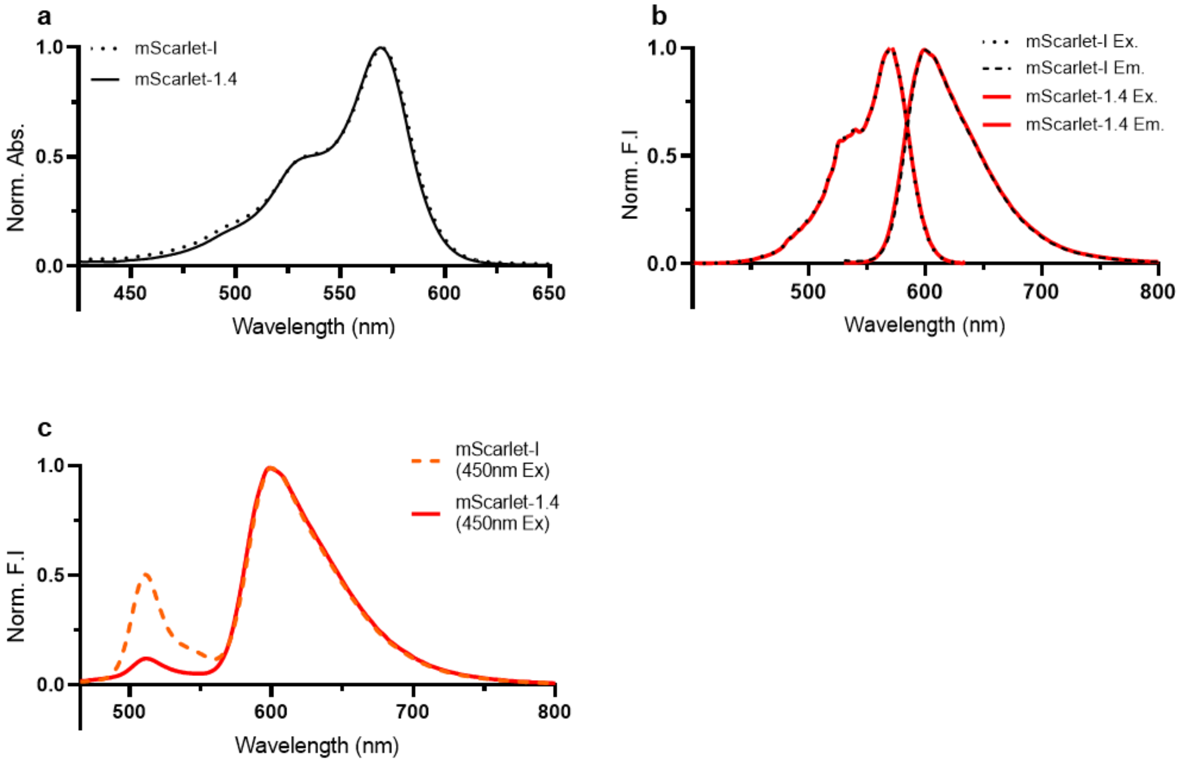


**Figure S2 Spectra of mScarlet-I1.4 and mScarlet-I. (a)** Normalized absorption spectra. **(b)** Normalized excitation and emission spectra (right). **(c)** Normalized emission spectra of mScarlet-I and mScarlet-I1.4 excited with 450nm (10nm bandwidth) light.

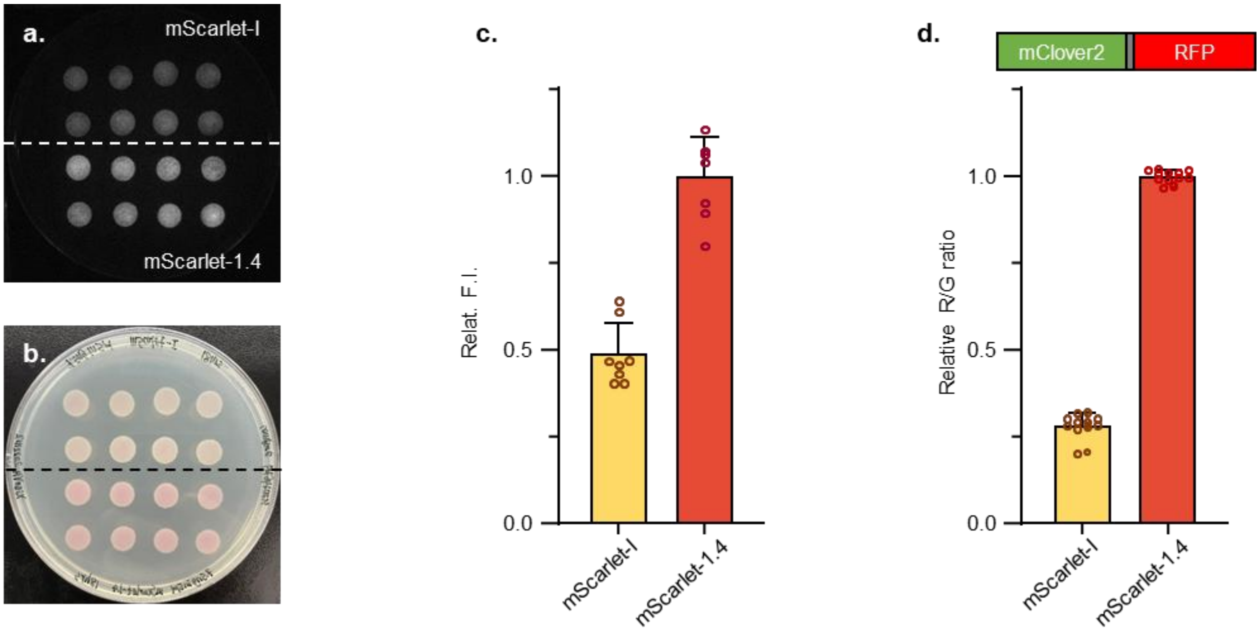


**Figure S3 Brightness comparison of mScarlet-I1.4 to mScarlet-I in E.coli. (a)** Red channel image of E.coli transfected with the same amount of pNCs_mScarlet-I or pNCs_mScarlet-I1.4 plasmids and cultured under 34 °C for 12 hr. **(b)** LED illuminated brightfield image of the same petri dish in **(a)**. **(c)** Relative red channel fluorescent intensity of E.coli, mScarlet-I1.4 is about 1 fold brighter than mScarlet-I, **(d)** Clover2 were fused to the N terminal of mScarlet-I and mScarlet-I1.4 with the same flexible linker, both of these fused FPs were cloned into pNCs vector and introduced into E.coli, after 12 hr culture under 34 °C in solid LB plate bacterial colonies were suspended in PBS and measured with microplate reader (Tecan M1000pro).

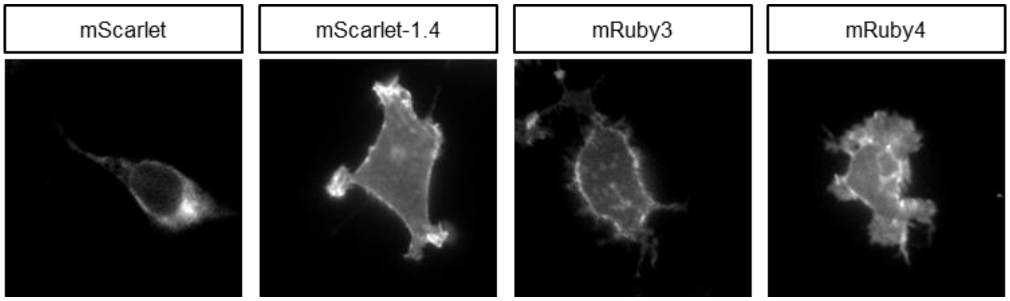


**Figure S4 Epifluorescence images showing expression and trafficking of extracellular loop1 (ECL1) inserted variants.** Scale bar = 20 μm.

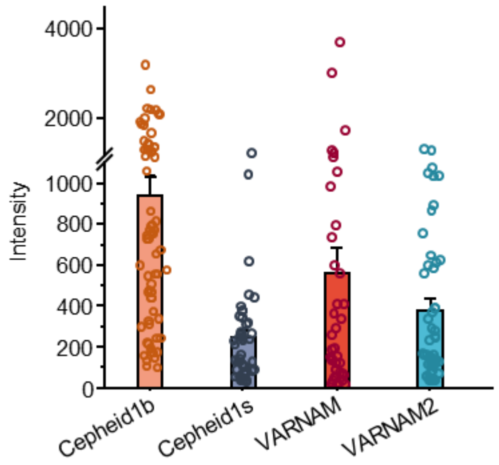


**Figure S5 Brightness comparison of Cepheid indicators with VARNAM family.** Each data point indicates the average photon count number in one cell, under wide field epifluorescence imaging with 561 nm laser.

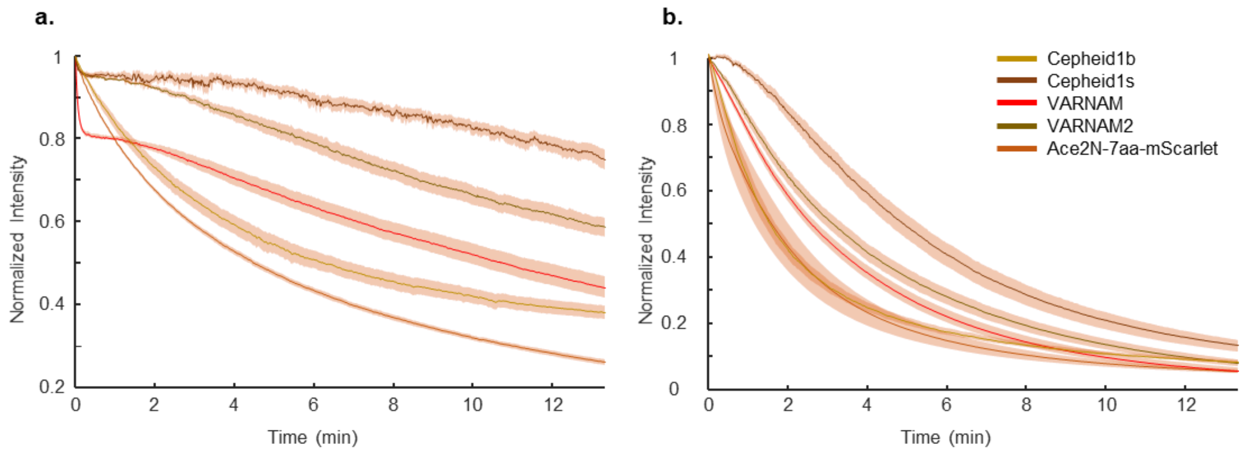


**Figure S6 Photo-stability comparison of Cepheid1 and other red GEVIs. (a)** 1.59 W/cm$^2$ 561 nm laser illumination (n = 7, 7, 6, 6, 5 cells). **(b)** 7.95 W/cm$^2$ 561 nm laser illumination (n = 7, 7, 7, 6, 5 cells);

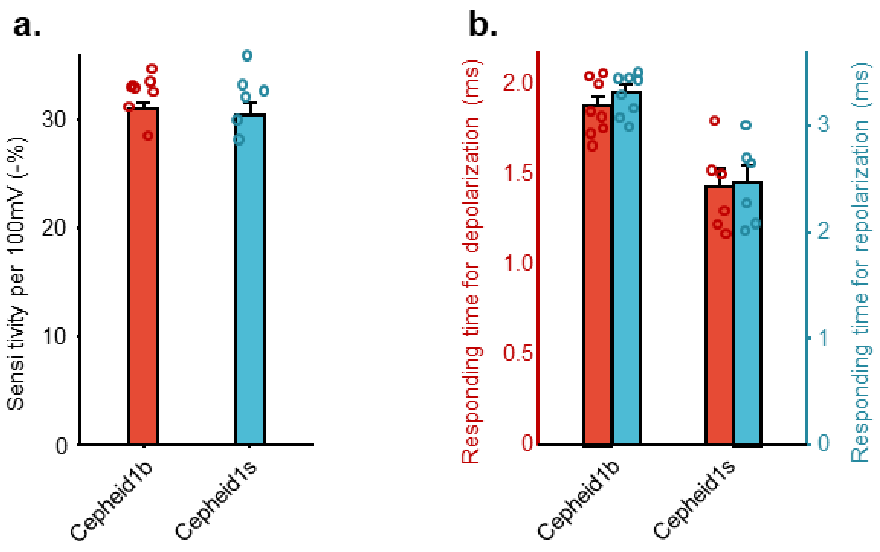


**Figure S7 Characteristics of Cepheid1 voltage response sensitivity and kinetics in HEK293T cells. (a)** sensitivity of Cepheid1b and Cepheid1s to 100 mV (-70 mV to +30 mV); **(b)** kinetics of Cepheid1b and Cepheid1s to depolarization (-70 mV to +30 mV) and to repolarization (+30 mV to -70 mV).

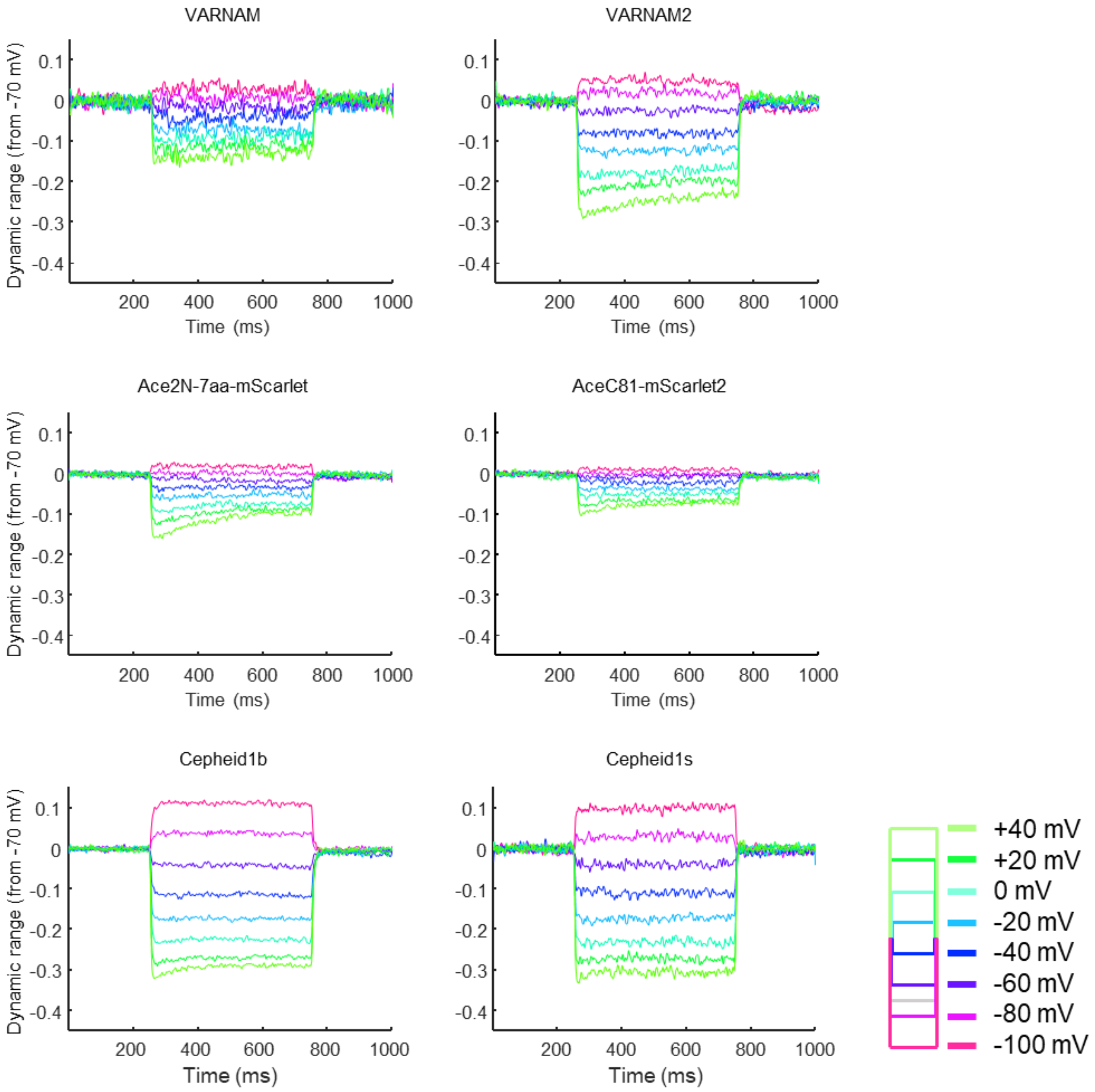


**Figure S8 Representative fluorescence traces of red GEVIs in response to a series of voltage steps.** The membrane potential was controlled via whole-cell voltage clamp, and a series of step waveforms were applied from −100 mV to 100 mV in increments of 20 mV increments). The dynamic range has been normalized to the fluorescence at membrane voltage $V_m$ = 70 mV.

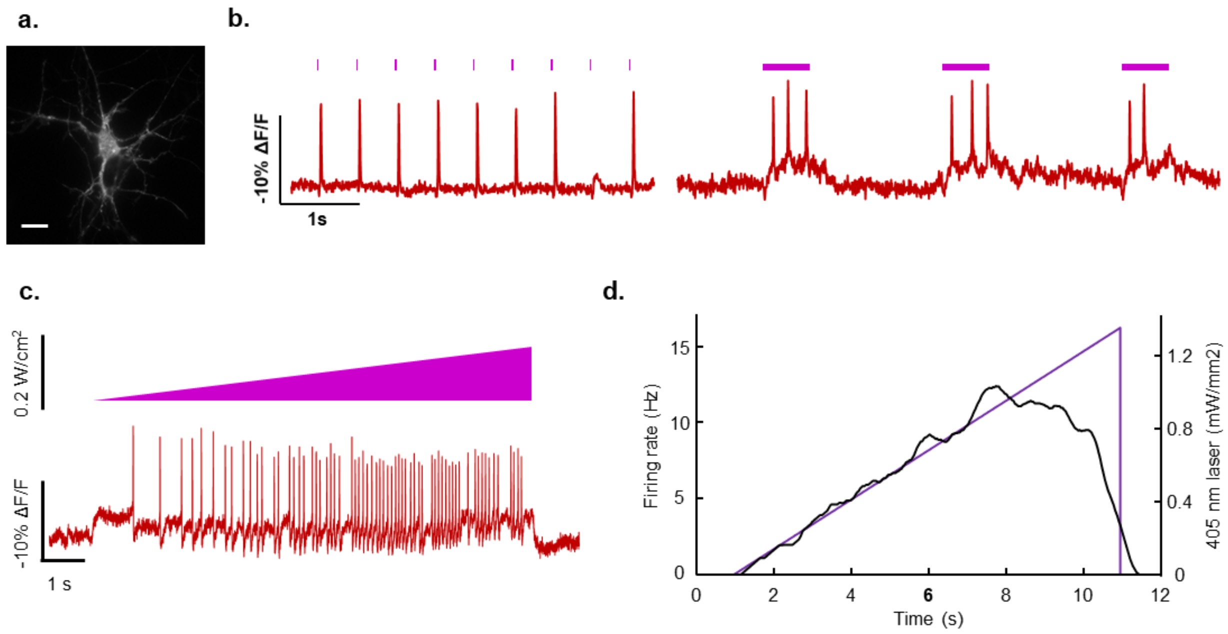


**Figure S9. All-optical electrophysiology in cultured neurons. (a)** Epifluorescence images showing expression of Cepheid1s-ST in a neuron transfected with Cepheid1s-ST-P2A-CheRiff (scale bar = 20 μm). **(b)** APs trigger by 405 nm pulse illumination, recorded by Cepheid1s-ST under wide field illumination with 561 nm laser. **(c)** APs trigger by 405 nm ramp illumination and recorded by Cepheid1s-ST. **(d)** Firing rate in the trial in **c** aligned with stimulation intensity.

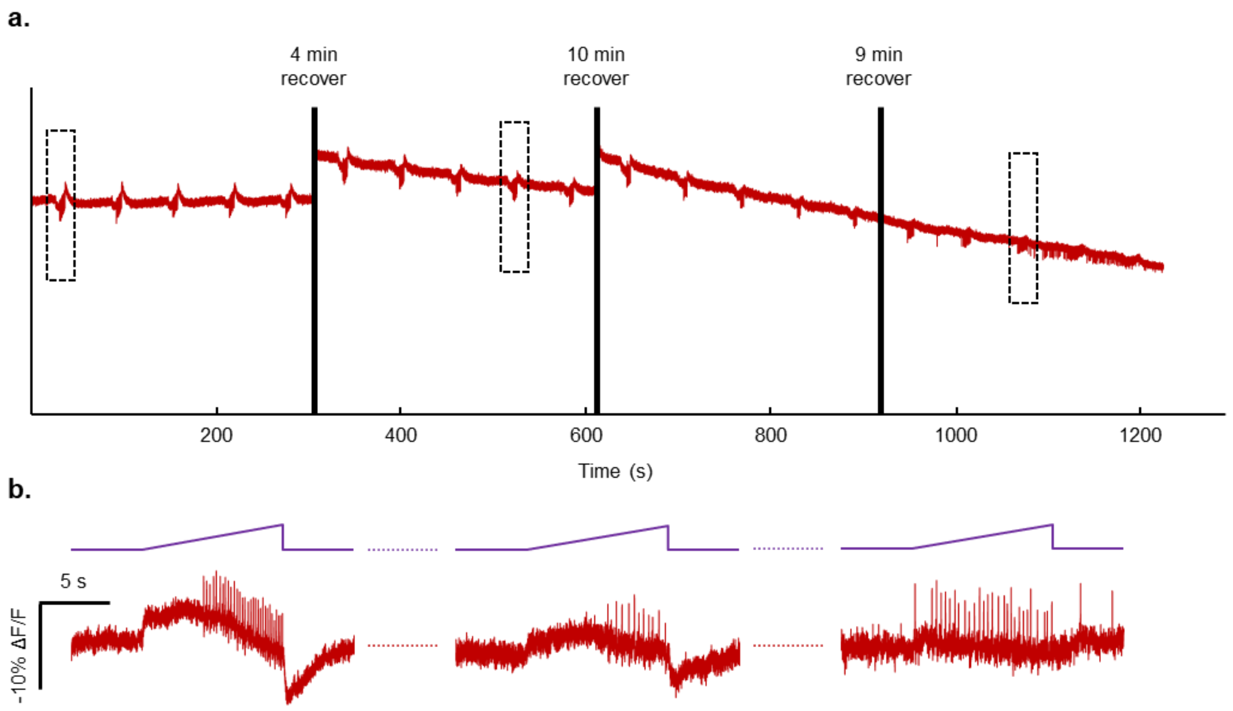


**Figure S10. long term optical recording under photo-activation of CheRiff. (a)** Photobleaching curve of the long-term imaging with Cepheid1s-ST. 10 s-long, ramp-shaped stimulation is conducted every 60s. The cultured neuron was recovered for 3-10 min in dark after a 300 s imaging. **(b)** Below, zoomed-in traces (red) of selected stimulation sessions in the dotted box in a, in alignment with optogenetic stimulation intensity patterns (magenta).

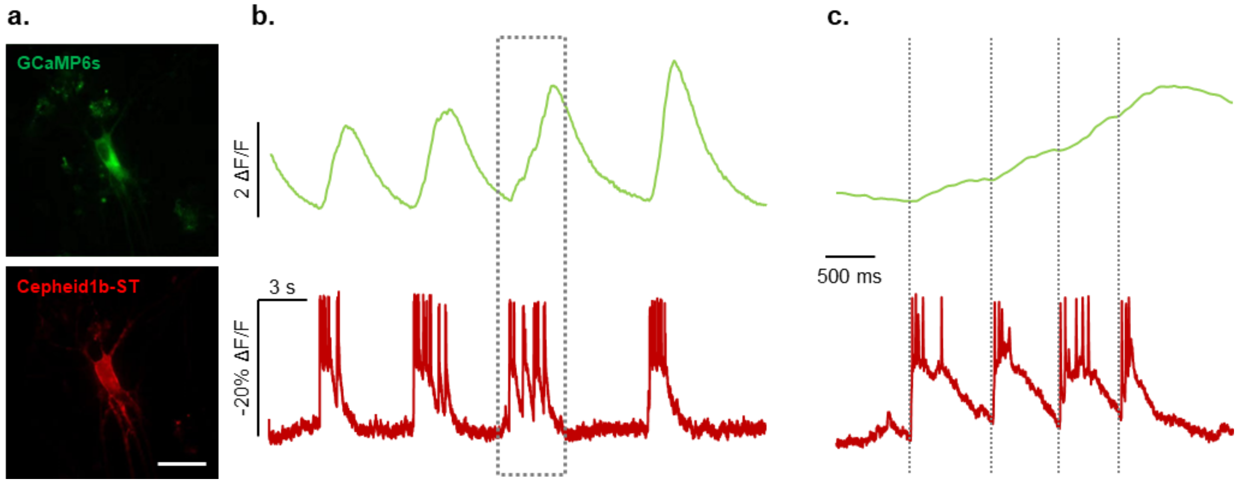


**Figure S11. Dual-color imaging using GCaMP6s and Cepheid1b-ST. (a)** Epifluorescence image of a neuron expressing GCaMP6s and Cepheid1b-ST. Scale bar, 20 μm. **(b)** Calcium and voltage activity in the same neuron. **(c)** A zoomed in view of fluorescence traces in the dashed gray box in **b**.

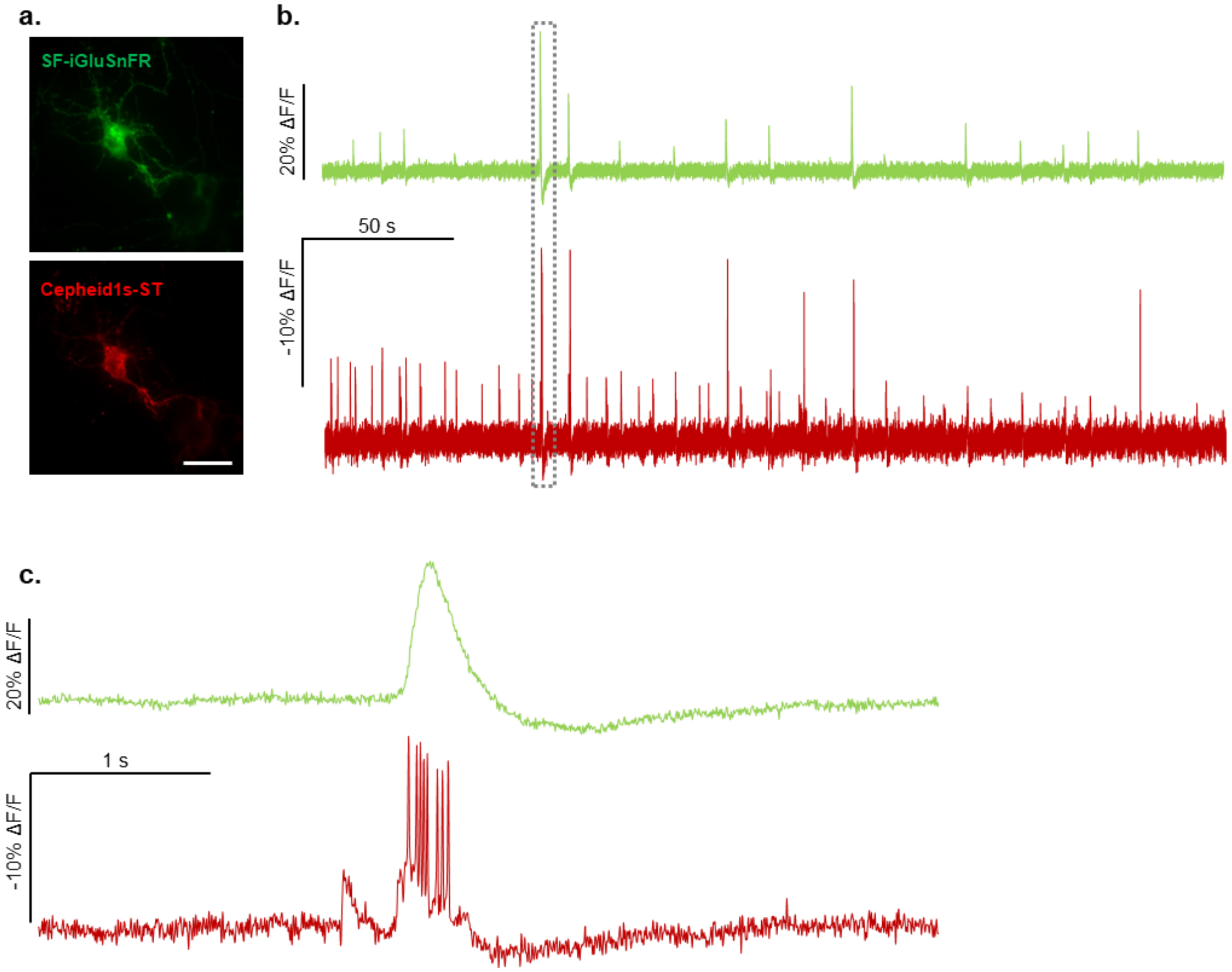


**Figure S12 Long-term optical recording of glutamate events and electrophysiology. (a)** Epifluorescence image of a neuron expressing SF-iGluSnFR and Cepheid1b-ST. Scale bar, 20 µm. **(b)** Glutamine and voltage activity in the same neuron. **(c)** A zoomed in view of fluorescence traces in the dashed gray box in **b**.
